## Supplementary information for "Young domestic chicks spontaneously represent the absence of objects"

### I. Length of trials

As we used a live setup, minor variance occurred in the length of trials. To rule out the potential effect of this variance, we encoded the length of trials and we investigated its effect on the dependent variables. First of all, we did not find significant difference between the length of trials in the two experimental conditions (Expected condition M = 31.34, SD = 1.81; Unexpected condition M = 31.15, SD = 1.89, Mann Whitney U test, $U$ = 22.56, $p$ = .35). Moreover, we did not find significant correlation either between the length of trials and Looking Time ($r(434) = .013$ , $p = .793$) or between the length of trials and Lateralization Index ($r(434) = -.003$ , $p$ = .949).

### II. Looking time data

| **Experiment** | **Condition** | **Total Mean ± SE** | **Male Mean ± SE** | **Female Mean ± SE** |
|---|---|---|---|---|
| Experiment 1 | Expected Disappearance | 12.67 ± 1.26 | 12.18 ± 1.48 | 13.2 ± 2.14 |
| | Unexpected Disappearance | 15.42 ± 1.41 | 14.27 ± 2.12 | 16.66 ± 1.87 |
| Experiment 2 | Expected Appearance | 14.56 ± 1.77 | 12.24 ± 2.85 | 15.57 ± 2.14 |
| | Unexpected Appearance | 12.91 ± 1.6 | 10.45 ± 2.14 | 15.03± 2.24 |
| Experiment 3 | Expected Appearance | 16.61 ± 1.59 | 16.08 ± 2.26 | 17.18 ± 2.3 |
| | Unexpected Appearance | 17.65 ± 1.55 | 18.42 ± 2.14 | 16.82 ± 2.3 |
| Experiment 4 | Expected Appearance | 13.03 ± 1.86 | 12.29 ± 2.6 | 15.27 ± 2.70 |
| | Unexpected Appearance | 13.55 ± 1.66 | 14.54 ± 2.56 | 12.60 ± 2.15 |

**Supplementary table 1**. **Average values of looking time data in Experiment 1-4**. We report the mean values and standard errors of looking times for the total sample size, and separately for males and females in the four experiments.
